# Identifying the source population of naturally recolonised rhinoceros (*Rhinoceros unicornis*) in Laokhowa-Burhachapori Wildlife Sanctuary, Assam, through multidisciplinary methods

**DOI:** 10.64898/2026.10.30.763177

**Authors:** Shrewshree Kumar, Jyoti Soun, Dibyendu Biswas, Megha Mehta, Jayanta Deka, Sonali Ghosh, Amit Sharma, Samrat Mondol

## Abstract

Megaherbivore population recovery largely depends on restoring connectivity between fragmented protected areas to facilitate natural dispersal, gene flow, and range expansion. The Indian rhinoceros (*Rhinoceros unicornis*) has recovered from severe population decline in the past, but several populations remain spatially restricted. Recent conservation efforts and the expansion of protected areas in Assam have coincided with natural rhino recolonization in Laokhowa-Burhachapori Wildlife Sanctuary (WLS). This has occurred from nearby regions of Orang NP and Kaziranga NP after the species was locally extinct in the 1980s. Identifying the source population of these recolonised rhinos is crucial for managing this emerging metapopulation. We used non-invasive genetic sampling, genetic assignment and a landscape connectivity model to investigate the origin and potential movement routes of the recolonising rhinos. We identified eight unique individuals from Laokhowa-Burhachapori WLS and assigned them to the potential source population using Bayesian and frequency-based approaches against a reference dataset of 365 individual rhinos across the rhino-bearing areas of Assam. Habitat suitability was modelled using 161 spatially thinned presence points and eight environmental variables. We applied least-cost path (LCP) analysis to assess landscape resistance between protected area nodes. All eight recolonising rhinos were assigned to Orang NP with high Bayesian assignment probabilities (q = 0.97-0.99). The connectivity model identified two least-cost pathways, with the Orang NP-Laokhowa-Burhachapori WLS linkage showing lower landscape friction than the Kaziranga NP linkage. These findings provide genetic evidence that the recolonisation occurred through natural dispersal, most likely from Orang NP, and indicate that recent landscape restoration has re-established the structural connectivity across this floodplain. Maintaining these linkages will be essential to promote natural range expansion and the long-term persistence of Indian rhinoceros in Assam.

## Introduction

Megaherbivores have experienced considerable range contractions and population declines since the Late Pleistocene, primarily due to human activities such as resource overexploitation, habitat fragmentation, and hunting (Morrison et al. 2007; Ripple et al. 2015; Smith et al. 2016; Malhi et al. 2016). The loss of megaherbivores has altered ecosystem functioning by reducing grazing, browsing, seed dispersal, nutrient recycling, and other ecosystem processes (Ripple et al. 2015; Malhi et al. 2016). Restoring megaherbivore populations has become a key conservation goal for maintaining landscape functionality and supporting species recovery (Trouwborst and Svenning 2022). Out of the at least fifty terrestrial megaherbivores that existed in the Late Pleistocene, only nine species are still extant (Malhi et al. 2016). Despite global efforts to recover their populations (Emsile and Brooks 1999; Madhusudan 2004; Brodie et al. 2011; Williams 2011), most megaherbivores are mainly found within protected areas (Trouwborst and Svenning 2022). However, because most protected areas worldwide are relatively small, they often cannot support viable populations of these large herbivores (Cantû-Salazar and Gaston 2010; Ripple et al. 2015). Therefore, expanding protected areas and maintaining connectivity between them are crucial strategies. These measures enable dispersal, promote genetic exchange, and are vital for the long-term survival of megaherbivore populations (Jhala et al. 2021; Brennan et al. 2022).

The Indian rhinoceros (*Rhinoceros unicornis*) is one of the remaining terrestrial megaherbivores (Malhi et al. 2016) endemic to the alluvial floodplains of the northern Indian subcontinent (Talukdar et al. 2025). Historically, this species ranged from the Indus Basin in present-day Pakistan to northeastern India and the Indo-Myanmar border (Rookmaker 1980; Rookmaker et al. 2016, 2024), but intense hunting and habitat destruction caused a 99.8% decline in its population and range during the early 20th century (Menon 1996; Mahmood et al. 2021). However, intensive conservation efforts, including anti-poaching strategies, habitat management, grassland restoration, and community engagement, have substantially increased the population, from approximately 200 individuals in the late 1900s to about 4000 in 2025 (Talukdar et al. 2025). Currently, the species inhabits about 20,000 sq.km. of fragmented protected areas across India and Nepal (Rookmaker 2004, 2016; Talukdar et al. 2008; Pant 2020). Within India, wild Indian rhinoceroses are primarily found in Uttar Pradesh, West Bengal, and Assam, with Assam hosting about 71% of the remaining population (Sharma 2022; Talukdar et al. 2025). In Uttar Pradesh and West Bengal, rhinos are confined to fragmented and isolated protected areas (PAs) (Mukherjee et al. 2020; Sharma 2022), which restricts opportunities for PA connectivity and genetic exchange (Ghosh et al. 2024). Conversely, several rhino-residing PAs in Assam are located within the floodplain system of the Brahmaputra river, connected by riverine grassland corridors that help maintain landscape connectivity (Borah et al. 2010; Jhala et al. 2020; Qureshi et al. 2023). Notably, the Kaziranga National Park (NP) and Orang NP complex function as essential floodplain corridors, supporting rhinos and a wide variety of other species (Borah et al. 2010; Qureshi et al. 2014). The Laokhowa-Burhachapori Wildlife Sanctuary (WLS), situated between Kaziranga National Park (NP) to the east and Orang NP to the west, is a strategically important part of this landscape. The sanctuary was home to approximately 41 wild rhinos in 1966, but widespread poaching and other disturbances during the Assam agitation in the 1980s led to the local extinction of the species here (Rookmaker 2024). Since 2007, there have been infrequent reports of stray rhinos, and two rhinos were translocated to the sanctuary in 2016 (Rookmaker 2024), but these efforts did not lead to a stable resident population. Recently, the Assam government has focused on expanding protected areas, with associated enhancement in habitat management and corridor protection within this landscape. These efforts resulted in the integration of approximately 200 sq. km. into the larger Kaziranga NP-Laokhowa-Burhachapori WLS-Orang NP complex, thereby improving ecological connectivity (Choudhury 2022; Ojah et al. 2023; Talukdar 2023). Since 2023, observations of natural rhino recolonisation in the Laokhowa-Burhachapori WLS (Talukdar et al. 2024) have begun, marking a rare instance of natural rhino movement driven by conservation efforts rather than human translocation. However, the origin of the recolonised rhinos and the migration pathways remain uncertain, with both Kaziranga NP and Orang NP being likely sources (Jhala et al. 2021). It is critical to ascertain these details to ensure the continued natural movement of more rhinos into this area, supporting the species’ natural range expansion.

In this study, we used robust, widely used genetic assignment tools to determine the source population of the recolonised rhinos and to identify potential dispersal pathways within the larger Kaziranga NP-Orang NP complex. In particular, we aimed to (i) identify the source of the newly migrated rhino individuals and (ii) evaluate whether the protected area boundary extension has facilitated any natural dispersal. We addressed these questions using 12 microsatellite loci surveyed in all newly arrived rhino individuals in Laokhowa-Burhachapori WLS, along with previously available genetic data from 335 rhinos (Ghosh et al. 2024) across all rhino-bearing areas in Assam. We believe that our results provide broader insights into conserving dispersal routes and the corridor functionality of these restored habitats.

## Materials and methods

### Research permission and ethical statements

The necessary field sampling permissions were accorded by the Assam Forest Department (Letter No. WL/FE.15/22(Pt-II)RhoDIS Workshop/2014 dated February 14, 2024) and Nagaon Wildlife Division (WD) (Letter No. WLN/B/Rhino General/2024/1824 dated November 28, 2024). As the entire study was based on non-invasive samples, no institutional ethical clearance was required.

### Study area

The study was carried out across all rhino-bearing protected areas in Assam, including Manas NP, Pobitora WLS, Orang NP, Kaziranga NP, and other regions. For this work, biological sampling was conducted in Laokhowa-Burhachapori WLS and Biswanath WD, while genetic data from other parks were obtained from (Ghosh et al. 2024).

Biswanath WD and Laokhowa-Burhachapori WLS are situated in the Brahmaputra floodplain of Assam, India. Biswanath WD covers 4.52 sq km in Biswanath district and is the tenth addition to Kaziranga NP (Assam Gazette No. FRM.203/2017/83). It lies on the northern bank of the Brahmaputra River (Sharma 2022), with its southern boundary adjoining Kaziranga NP (Hazarika 2020). Laokhowa WLS covers an area of 70.1 sq km in Nagaon district, while Burhachapori WLS covers an area of 44.06 sq km in Sonitpur district (Ojah et al. 2015). The two sanctuaries are separated by Dhania Suti, a channel of the Brahmaputra River. Together, they form a contiguous protected landscape, Laokhowa-Burhachapori WLS (Ojah et al. 2023). Laokhowa-Burhachapori WLS lies between Kaziranga NP to the east and Orang NP to the west, serving as an important migratory corridor between these two major protected areas (Ojah et al. 2023; Talukdar et al. 2024). This riverine landscape is a highly fertile habitat with extensive grasslands (*Saccharum spp.*, *Arundo spp.*, *Erianthus sp.*, *Phragmites spp.*, etc.), interrupted by mixed woodlands (*Bombax sp.*, *Albizzia spp.*, *Careya sp.*, *Lagerstroemia sp.*, *Zizyphus sp.*, etc.), freshwater mangroves (*Barringtonia sp.*, *Syzyzium sp.*, etc.), and wetlands (Ojah et al. 2023). This area supports a rich floral and faunal diversity, including many critically endangered grassland-dependent species such as the Bengal florican (*Houbaropsis bengalensis*), Greater adjutant stork (*Leptoptilos dubius*), Black-bellied tern (*Sterna acuticada*), Asiatic wild buffalo (*Bubalus arnee*), Hispid hare (*Caprolagus hispidus*), and Eastern swamp deer (*Rucervus duvaucelii ranjitsinhi*). In 2023, a 195 sq. km area was added as the first extension to Burhachapori WLS, stretching from the western edge of Kaziranga NP to the eastern boundary of Orang NP (Assam Gazette Ecf No. 274705/23). Earlier, a further 200 sq. km. was extended from Orang NP and designated as the park’s second addition in 2022, neighbouring the first extension of Burhachapori WLS (Assam Gazette No. FRW.14/2004/Pt/64), enhancing landscape connectivity across the Kaziranga-Orang larger landscape, hereafter, the Kaziranga Orang Riverine Landscape (Borah et al. 2010). Following habitat expansion, the Laokhowa-Burhachapori WLS in Assam is currently considered a suitable habitat for rhinos (Ojah et al. 2015), and recent natural dispersal of Indian rhinos to this park has been documented (Talukdar et al. 2024; Melletti et al. 2025). Given its geographical proximity to Orang NP and Kaziranga NP (Jhala et al. 2021), dispersing individuals are likely to originate from either of these source populations.

### Biological sampling

We planned a genetic assignment approach to identify the source population of migrating rhinos in the Laokhowa-Burhachapori WLS. This was accomplished by collecting fresh rhino dung samples from the park and comparing them with available genetic data from rhinos across different parks in Assam. Indian rhinos typically defecate in 6-7 boluses at a time, at fixed locations known as dung heaps or middens (Laurie 1982; Hazarika and Saikia 2010). For this study, we conducted intensive field surveys (using foot, boat, and vehicle as appropriate to local conditions) in March and December 2024 to identify midden sites (Figure 1a). Combining both sampling efforts, we collected nine fresh dung samples from the entire area. Additionally, we collected 33 fresh dung samples from the Biswanath WD of Kaziranga NP in March 2024. Once the dung piles were identified, sampling was carried out according to the already standardized approach (Ghosh et al. 2022). In brief, the uppermost mucus layer of the fresh dung bolus was swabbed in duplicate with sterile swabs (Himedia, Mumbai, India) and the swabs were stored in 2 ml sterile tubes. The geo-tagged samples were transported to the laboratory at the Wildlife Institute of India and stored in a -20°C freezer until further processing.

**Figure 1:**
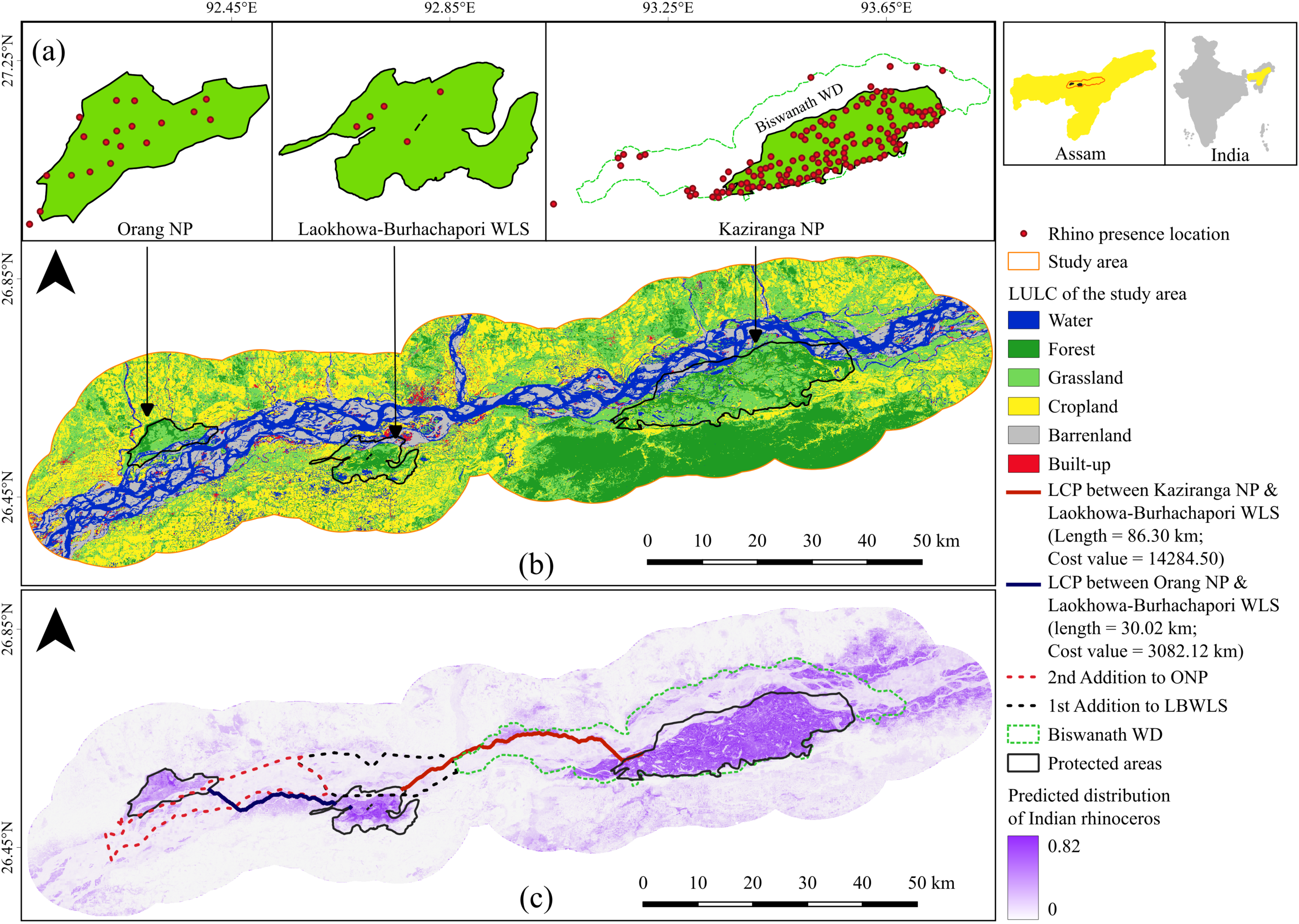
Study area and landscape connectivity modelling in the Orang NP-Laokhowa-Burhachapori WLS-Kaziranga NP landscape in Assam, India. (a) Rhino dung sampling/presence locations used for habitat suitability modelling in the respective parks. (b) Land-use/land-cover map in the Orang NP-Kaziranga NP complex. (c) Predicted distribution of the Indian rhinoceros and least-cost pathways, together with protected areas and their additions. The lengths of the least-cost pathways and cost values are provided in the legend.

### DNA extraction, individual identification, molecular sexing, and genetic relatedness

Genomic DNA was extracted from the swabs following the standardized protocol for Indian rhinos described in Ghosh et al. (2022). In brief, swabs were digested overnight at 56 °C in ATL (Animal Tissue Lysis buffer) and Proteinase K (20 mg/ml), followed by extraction using the modified QIAamp DNA Tissue Kit (QIAGEN Inc., Hilden, Germany) extraction protocol outlined in Ghosh et al. (2022). DNA was eluted twice into 100 μl of preheated (70°C) 1X TE buffer and stored at -20°C until further use. Extraction negatives were included in each set (n=23) of the experiment to monitor potential contamination.

Unique individual rhinos were identified using a panel of 12 previously standardized microsatellite loci (Ghosh et al. 2021). Sex identification was performed on genetically identified individuals, following the protocol of Kumar et al. (2025). For both, PCRs were carried out in 10 μL reaction volumes containing 4 μL of 2X Qiagen multiplex PCR buffer mix (QIAGEN Inc., Hilden, Germany), 0.25 μM primer mix, 4 μM BSA (4 mg/mL), and 2 μL of template DNA. The thermal cycling conditions included an initial denaturation (95°C for 15 min); 40 cycles consisting of denaturation (95°C for 30 s), annealing (Ta °C for 40 s), and extension (72°C for 40 s); followed by a final extension (72°C for 30 min). Each reaction set included PCR and extraction negative controls (to monitor contamination), along with a reference rhino DNA sample as a positive control. Each sample was genotyped thrice using the LIZ 500 size standard (Applied Biosystems, California, USA) in an ABI 3500XL Genetic Analyzer (Applied Biosystems, California, USA) and scored manually with GENEMARKER (Softgenetics Inc., Pennsylvania, USA) following established Indian rhino allele bins as described in Ghosh et al. (2021) and Kumar et al. (2025). Unique multilocus genotypes were identified using CERVUS (Kalinowski et al. 2007). The summary statistics, including the probability of identity for both siblings (PIDsibs) and unrelated individuals (PIDunbiased), as well as observed heterozygosity (Ho), were calculated in GIMLET (Valiere 2002).

To determine possible parent-offspring and sibling relationships, pairwise relatedness among individuals was calculated using the program ML-RELATE (Kalinowski et al. 2006) with a maximum likelihood estimation approach. Four relationship categories (parent-offspring (PO), full siblings (FS), half siblings (HS), and unrelated (U)) were assessed with 1000 permutations at a 95% confidence level.

### Genetic assignment of unknown rhino individuals

To identify the source populations of the dispersed rhinos, we employed both Bayesian and frequency-based genetic assignment methods. For this, we used the genetic data generated in this study (Laokhowa-Burhachapori WLS and Biswanath WD) alongside previously available data (Ghosh et al. 2024) from 335 rhinos across four different rhino-bearing areas (Kaziranga NP (n = 163), Orang NP (n = 55), Manas NP (n = 26), and Pobitora WLS (n = 91)).

The assignment analyses were carried out by initially assigning the Laokhowa-Burhachapori WLS individuals to all four Assam rhino populations (Kaziranga NP, Manas NP, Orang NP, and Pobitora WLS) to assess their general population affinity, followed by specific assignments to the target populations (Kaziranga and Orang NPs). For the Bayesian approach, the protocol of Ghosh et al. (2024) was employed using the USEPOPINFO model in STRUCTURE (Pritchard et al. 2000), with updated allele frequencies and no admixture assumed. The number of genetic clusters was set at the previously determined K = 3 (Ghosh et al. 2024). The POPFLAG option was utilized to designate reference individuals and to assign unknown genotypes to predefined populations, with a threshold of q > 0.95 for confident assignment, as recommended for the Assam rhino population (Ghosh et al. 2024). Frequency-based assignment was performed in ARLEQUIN v3.1 (Excoffier et al. 2005) using the genotype assignment test, which estimates the log-likelihood of each individual’s multilocus genotype originating from each reference population (Paetkau et al. 2004; Waser and Strobeck 1998).

### Development of a habitat permeability layer and least-cost path

Recent government-led PA expansion and corridor protection efforts are likely the primary drivers of natural rhino movement within the contiguous Kaziranga Orang Riverine Landscape. To identify the critical corridors and habitat patches for targeted conservation efforts, the ‘least-cost path’ modelling approach was used. The study area was delineated as a polygon with a 10 km buffer around the perimeter of the three protected areas (Kaziranga NP, Laokhowa-Burhachapori WLS, and Orang NP) within the Kaziranga-Orang complex. The buffer was intended to include potential dispersal routes between the three protected areas, regarded as focal nodes, and to cover the wider landscape relevant to potential rhino movements.

The least cost path (LCP) analysis using ‘leastcostpath’ package (version 2.0.13) in R requires two main components: i) the core area polygons or nodal points between which the connectivity is to be estimated; ii) a landscape resistance layer indicating the relative ease or difficulty for the movement of focal species (Lewis 2021). The GIS layers for the protected areas were used to define the core area polygon. The resistance layer was developed using an ensemble species distribution modelling (SDM) approach implemented in R (Version 26.06.0) using ‘sdm’ package (Naimi and Araújo 2016). A total of n=656 GPS locations of rhino dung samples (from the present study and Ghosh et al. 2024) served as presence points for model building and calibration. These locations represent confirmed rhino presence and thus indicate relevant potential structural connectivity. Furthermore, spatial thinning of the presence points was conducted using the ‘spThin’ package in R (Aiello-Lammens et al. 2015) to minimize spatial autocorrelation caused by clustered points. After thinning, n=161 presence points were retained, with a minimum distance of 1 km between points. Based on the species’ biology and the study area size (Kaziranga NP-Orang NP complex), we selected eight environmental variables as predictors: Land-use and land-cover (LULC) (Source: Developed in Google Earth Engine using USGS/NASA Landsat 8 and Landsat 9 Collection 2, Tier 1 Surface Reflectance (L2) imagery at a 30-meter spatial resolution with multi-season composite approach), Normalised Difference Vegetation Index (NDVI) (Source: Developed in Google Earth Engine using Landsat 8-9 OLI/TIRS C2 L2 imagery), Human Footprint (HFP) (Source: Gassert et al., 2023), Forest Cover Type (FC) (Source: Forest Survey of India), Elevation (ELE) (Source: NASA JPL. (2020), Slope (SLP) (Source: Developed from Digital Elevation Model layer), Terrain Ruggedness Index (TRI) (Source: Developed from Digital Elevation Model layer), and Nightlight (NTL) (NOAA VIIRS Day/Night Band (DNB) Composite).

Prior to the distribution modelling, variance inflation factor (VIF) stepwise (vifstep) analysis was executed to diagnose multicollinearity among the predictor layers (Naimi et al. 2014). Three modelling algorithms were fitted using the ‘sdm’ package, namely Random Forest (RF), Boosted Regression Tree (BRT), and Generalised Linear Modelling (GLM) (Breiman 2001; Friedman 2001; McCullagh and Nelder 1989; Naimi and Araújo 2016). 1000 background points were generated following Barbet-Massin et al. (2012) to represent the environmental space of the study area, excluding known presence locations. Each model was evaluated through a random sub-sampling cross-validation approach, with a 70:30 train-test split and 10 replicates. For better model prediction, only the models with a mean area under the curve of ‘receiver operating characteristic’ (ROC-AUC) ≥ 0.85 were retained and ensembled (Araújo and New 2007; Swets 1988; Fielding and Bell 1997). The output was a raster showing rhino habitat suitability, with higher values indicating greater suitability and lower values indicating less suitability. This suitability surface was used to represent landscape permeability and was subsequently transformed by taking its reciprocal to produce a resistance layer, with higher values representing increased movement costs. Further, the structural connectivity was modelled across the study area by establishing a spatial network connecting the three focal PAs or nodes, which are the polygons of Kaziranga NP, Laokhowa-Burhachapori WLS and Orang NP. The LCP was constructed connecting the centroids of Kaziranga NP, Laokhowa-Burhachapori WLS and Orang NP. This was followed by the extraction of the LCPs only for the outside PA boundaries along with associated ‘accumulated cost value’ (indicator of absolute landscape friction). This cost was calculated from the cumulative sum of resistance values from the pixels along the single-pixel paths generated.

## Results

### Individual and sex identification

Out of 42 faecal samples collected (33 from Biswanath WD and nine from Laokhowa-Burhachapori WLS), 38 individual animals were identified (n=30 from Biswanath WD and n=8 from Laokhowa-Burhachapori WLS). In total, four recaptures were identified in the dataset. After including these individuals from both areas, the overall number of rhinos in the Assam dataset reached 373.

The sex identification of the individual animals revealed 12 males and 15 females from Biswanath WD (three samples failed to produce data), and seven males and one female from Laokhowa-Burhachapori WLS. The cumulative PID_(sibs)_ and PID_(unbiased)_ values were 1.55 x 10^-3^ and 2.93 X 10^-7^ for Biswanath WD, and 3.86 X 10^-4^ and 1.57 X 10^-8^ for Laokhowa-Burhachapori WLS. The summary statistics for the new areas are provided in Table 1. The genetic relatedness analyses revealed five pairs of full-siblings (FS) among individuals in Biswanath WD and one FS relationship among individuals in Laokhowa-Burhachapori WLS. ***Assignment of unknown individuals from Laokhowa-Burhachapori WLS***

**Table 1:** Summary statistics of microsatellite genotyping results for Indian rhinoceros used in this study from Biswanath WLD and Laokhowa-Burhachapori WLS in Assam, India. He=expected heterozygosity, Ho=observed heterozygosity, PIDsibs=probability of identity among siblings and PIDunbiased=unbiased probability of identity.

| Population | Loci | Number of alleles | Samples collected | Individuals identified | He | Ho | PIDsibs | PIDunbiased |
|---|---|---|---|---|---|---|---|---|
| **Biswanath WD** | RH1 | 3 | 33 | 30 | 0.62 | 0.33 | 1.55 x 10-3 | 2.93 X 10-7 |
|  | RH3 | 3 |  |  | 0.59 | 0.27 |  |  |
|  | RH10 | 5 |  |  | 0.69 | 0.5 |  |  |
|  | IR10 | 1 |  |  | 0 | 0 |  |  |
|  | RH7 | 2 |  |  | 0.31 | 0.1 |  |  |
|  | SR63 | 2 |  |  | 0.49 | 0.03 |  |  |
|  | RH5 | 4 |  |  | 0.53 | 0.07 |  |  |
|  | IR12 | 2 |  |  | 0.5 | 0 |  |  |
|  | 7B | 2 |  |  | 0.22 | 0 |  |  |
|  | RH9 | 3 |  |  | 0.4 | 0.13 |  |  |
|  | RH4 | 5 |  |  | 0.76 | 0.4 |  |  |
|  | RH11 | 4 |  |  | 0.65 | 0.27 |  |  |
|  | **Mean** | **3 ± 1.27** | **33** | **30** | **0.48 ± 0.21** | **0.17 ± 0.17** | **1.55 x 10<sup>-3</sup>** | **2.93 X 10<sup>-7</sup>** |
| **Laokhowa-Burhachapori WLS** | RH1 | 3 | 9 | 8 | 0.53 | 0.25 | 3.86 X 10-4 | 1.57 X 10-8 |
|  | RH3 | 2 |  |  | 0.47 | 0 |  |  |
|  | RH10 | 2 |  |  | 0.42 | 0.13 |  |  |
|  | IR10 | 2 |  |  | 0.43 | 0.13 |  |  |
|  | RH7 | 2 |  |  | 0.46 | 0.13 |  |  |
|  | SR63 | 2 |  |  | 0.44 | 0 |  |  |
|  | RH5 | 2 |  |  | 0.46 | 0.38 |  |  |
|  | IR12 | 0 |  |  | 1 | 0 |  |  |
|  | 7B | 0 |  |  | 1 | 0 |  |  |
|  | RH9 | 2 |  |  | 0.46 | 0.13 |  |  |
|  | RH4 | 3 |  |  | 0.63 | 0.25 |  |  |
|  | RH11 | 2 |  |  | 0.5 | 0.38 |  |  |
|  | **Mean** | **1.8 ± 0.93** | **9** | **8** | **0.56 ± 0.21** | **0.14 ± 0.14** | **3.86 X 10<sup>-4</sup>** | **1.57 X 10<sup>-8</sup>** |

In the Bayesian assignment analyses with four potential source populations (Kaziranga NP, Manas NP, Orang NP, and Pobitora WLS), the eight rhinos were assigned to Orang NP, with strong assignment probabilities ranging from q = 0.978 - 0.996 (Table 2a). However, the frequency-based method assigned six individuals to Orang NP with log-likelihood values from -21.537 to -31.902, while two individuals were assigned to Pobitora WLS (−26.830 and - 28.124) (Table 2b). Only Kaziranga and Orang NP-driven Bayesian assignment outcomes also assigned all eight individuals to Orang NP (q = 0.980-0.996) (Table 2a). The frequency-based method assigned seven individuals to Orang NP with log-likelihood values ranging from - 21.537 to -29.384, and one individual to Kaziranga NP (−27.717) (Table 2b). Together, we ascertained that all eight rhinos migrated from Orang NP to Laokhowa-Burhachapori WLS. ***Landscape connectivity and least-cost corridor pathways***

**Table 2:** Genetic assignment of Indian rhinoceros individuals identified in Laokhowa-Burhachapori WLS in Assam, India, to potential source populations. (a) Bayesian assignment probabilities (q) based on analyses including all three reference populations (Kaziranga NP, Orang NP and Pobitora WLS) and analyses restricted to Orang NP and Kaziranga NP. (b) Frequency-based assignment log-likelihood values for the same reference-population sets.

| (a) |  |  |  |  |  |  |
| **Bayesian assignment (using all three population)** |  |  | **Bayesian assignment (using Orang NP & Kaziranga NP)** |  |  |  |
| **Individuals** | **q value** | **Assignment** | **Individuals** | **q value** | **Assignment** |  |
|---|---|---|---|---|---|---|
| LBWLS_Ind1 | 0.995 | All assigned to ORANG | LBWLS_Ind1 | 0.995 | All assigned to ORANG |  |
| LBWLS_Ind2 | 0.996 |  | LBWLS_Ind2 | 0.996 |  |  |
| LBWLS_Ind3 | 0.982 |  | LBWLS_Ind3 | 0.982 |  |  |
| LBWLS_Ind4 | 0.980 |  | LBWLS_Ind4 | 0.980 |  |  |
| LBWLS_Ind5 | 0.978 |  | LBWLS_Ind5 | 0.981 |  |  |
| LBWLS_Ind6 | 0.992 |  | LBWLS_Ind6 | 0.992 |  |  |
| LBWLS_Ind7 | 0.986 |  | LBWLS_Ind7 | 0.986 |  |  |
| LBWLS_Ind8 | 0.995 |  | LBWLS_Ind8 | 0.995 |  |  |
| (b) |  |  |  |  |  |  |
| **Frequency-based assignment (using all three population)** |  |  |  | **Frequency-based assignment (using Orang NP & Kaziranga NP)** |  |  |
| **Individuals** | **Kaziranga NP** | **Orang NP** | **Pobitora WLS** | **Individuals** | **Kaziranga NP** | **Orang NP** |
| LBWLS_Ind1 | -27.622 | **-21.537** | -29.189 | LBWLS_Ind1 | -27.622 | **-21.537** |
| LBWLS_Ind2 | -29.550 | **-22.611** | -24.966 | LBWLS_Ind2 | -29.550 | **-22.611** |
| LBWLS_Ind3 | -26.633 | **-24.517** | -27.721 | LBWLS_Ind3 | -26.633 | **-24.517** |
| LBWLS_Ind4 | -26.617 | **-25.496** | -27.701 | LBWLS_Ind4 | -26.617 | **-25.496** |
| LBWLS_Ind5 | -27.717 | -29.441 | **-26.830** | LBWLS_Ind5 | **-27.717** | -29.441 |
| LBWLS_Ind6 | -36.619 | -29.384 | **-28.124** | LBWLS_Ind6 | -36.619 | **-29.384** |
| LBWLS_Ind7 | -32.024 | **-27.851** | -34.421 | LBWLS_Ind7 | -32.024 | **-27.851** |
| LBWLS_Ind8 | -44.533 | **-31.902** | -45.241 | LBWLS_Ind8 | -44.533 | **-31.902** |

None of the eight environmental variables showed correlation and was therefore retained for further modelling Table 3. The performance metrics for RF, BRT, and GLM were statistically significant, with test AUC values of 0.91 for RF and 0.89 for both GLM and BRT, exceeding the 0.80 threshold for ensembling (Figure 2a). Using AUC metrics, ’variable importance’ identified NTL, NDVI, and ELE as the top predictors at 28.5%, followed by HFP at 23.4% (Figure 2b). Moreover, the response curve indicates a sharp decrease in distribution as NTL and ELE increase, while the distribution rises with higher NDVI values (Figure 2c), indicating that human activities, vegetation condition, and ground elevation are the primary factors influencing habitat suitability or permeability in this area. The combined map reveals that the most suitable zones for rhinos are primarily situated within or close to protected areas. In contrast, suitability diminishes in regions that have been heavily altered by human actions, such as agricultural landscapes and higher elevation zones (Figure 2c).

**Figure 2:**
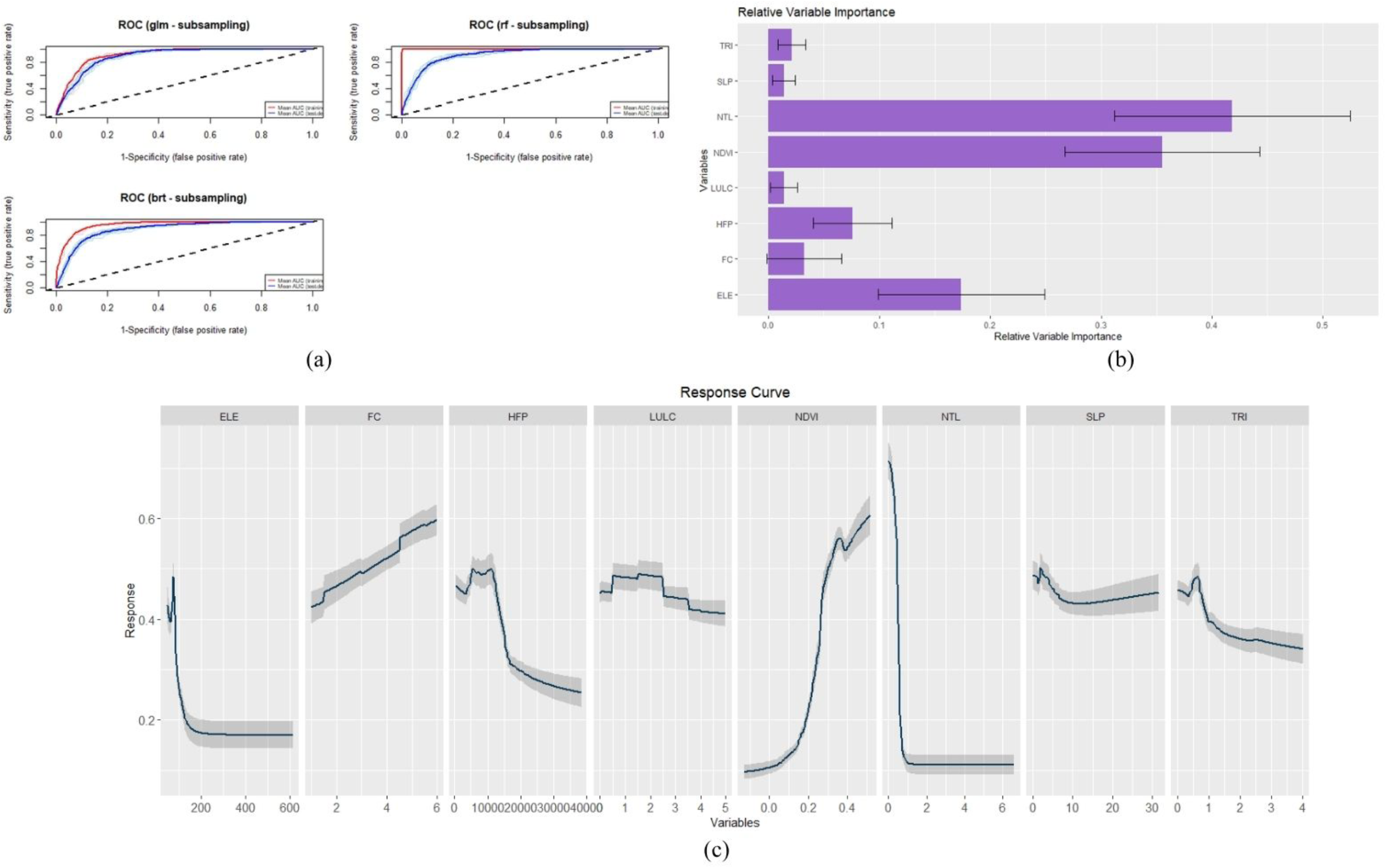
Performance and variable contributions of the species distribution model for the Indian rhinoceros in the Orang NP-Laokhowa-Burhachapori WLS-Kaziranga NP landscape in Assam, India. (a) Receiver operating characteristic (ROC) curves for the three modelling algorithms-Generalised Linear Model (GLM), Random Forest (RF), and Boosted Regression Tree (BRT), evaluated using random subsampling. (b) Relative importance of the eight environmental variables included in the ensemble modelling: Terrain Ruggedness Index (TRI), Slope (SLP), Nighttime lights (NTL), Normalised Difference Vegetation Index (NDVI), Land-Use/Land-Cover (LULC), Human Footprint Index (HFP), Forest Cover (FC), and Elevation (ELE). (c) Response curves showing the relationship between each environmental variable and predicted Indian rhinoceros distribution.

**Table 3:** Variance Inflation Factor (VIF) values for environmental predictor variables used in this study for species distribution model for the Indian rhinoceros in the Kaziranga NP-Orang NP landscape in Assam, India

| **Variable** | **Variable Inflation Factor (VIF)** |
|---|---|
| Slope (SLP) | 2.64 |
| Terrain Ruggedness Index (TRI) | 2.02 |
| Elevation (ELE) | 2.06 |
| Forest Cover (FC) | 2 |
| Human Footprint Index (HFI) | 1.66 |
| Normalized Difference Vegetation Index (NDVI) | 1.44 |
| Nighttime Lights (NTL) | 1.39 |
| Land Use Land Cover (LULC) | 1.12 |

The analyses identified two major LCPs: one between Kaziranga NP and Laokhowa-Burhachapori WLS, measuring 86.30 km with a cost value of 14284.50, and the other between Orang NP and Laokhowa-Burhachapori WLS, spanning 30.02 km with a cost of 3082.12 (Figure 1c).

## Discussion

The recent natural recolonization of rhinos in Laokhowa-Burhachapori WLS marks a significant conservation milestone for the Indian rhinoceros in Assam, following their local extinction over the past few decades (Talukdar 2024; Rookmaker 2024). Although the natural migration event was well established (Talukdar 2024), the integrated genetic and landscape modelling approach confirmed Orang NP as the source population and identified critical areas between Orang NP and Kaziranga NP that require focused, long-term conservation attention. The Bayesian assignment method confidently placed all individuals in Orang NP, with 97%-99% certainty, whereas the frequency-based approach yielded mostly consistent results, assigning six individuals to Orang NP and two to Pobitora WLS. These differences likely stem from the low genetic differentiation among the Assam rhino populations (F_ST_ = 0.03 between Orang NP and Pobitora WLS), as noted by Ghosh et al. (2024). Frequency-based methods have limited ability to distinguish weakly differentiated populations because they rely on likelihood estimates from predefined reference groups. Conversely, Bayesian clustering methods incorporate allele frequency data and tend to perform better when genetic differences are small (Latch 2006). The inconsistent assignment of two individuals to Pobitora WLS might also be due to missing data in two samples, which could reduce assignment accuracy, especially when genetic variation among reference populations is minimal. The data analyses were based on one of the most extensive genetic database of the Indian rhinoceros in Assam, with new data from Biswanath WD and Laokhowa-Burhachapori WLS added to earlier data (Ghosh et al. 2024). Such effort also demonstrates the utility of a genetic database in addressing management-focused challenges for other habitat-specialist species, as well as other applications (for example, understanding population genetic patterns (Gaggiotti et al. 2009; Qiao et al. 2019; Klingler et al. 2021; Kolipakam et al. 2019; Ghosh et al. 2024), assessing local-scale genetic connectivity (Sawaya et al. 2014; Marrotte et al. 2017; Biswas et al. 2022; Tyagi et al. 2024), forensic investigations (Millions et al. 2006; Harper et al. 2018; Wasser et al. 2018) and identifying source populations for translocations (Glowatzki-Mullis et al. 2008; Weeks et al. 2017; Coleman et al. 2013).

Our molecular sexing data suggest a strong male-biased composition of the recolonizing population, as seven of the eight identified individuals were male. Such sex-biased dispersal patterns have also been documented in Black rhino (Mellya et al. 2025), Alashan red deer (Gao et al. 2025), Norwegian red deer (Haanes et al. 2011) and wild boar (Pérez-González et al. 2025). Although the current sample size is small, the observed sex ratio may indicate that males initially drive exploratory dispersal into newly available habitats (Laurie 1982). However, population viability analyses by Jhala et al. (2021) modelled a female-biased founder population for a newly established rhino population, highlighting the importance of female recruitment for long-term persistence. If natural female movement remains limited, introducing new females in the area may be necessary to support the establishment of a healthy, resident rhino population. A similar strategy to that implemented in Manas NP should be followed, where mixing of different rhino gene pools from various sites within Assam can be pursued. As the recolonizing individuals were assigned from Orang NP, other rhino populations in Assam (Pobitora WLS, Kaziranga NP) should be considered the sources of females for translocation (Ghosh et al. 2024). Some of the migrated individuals were observed outside the Laokhowa-Burhachapori WLS park boundary, near the first addition area of Burhachapori WLS, close to Kaziranga NP. These animals are likely exploring this region before settling into their territories. While identifying the exact movement pathways requires more intensive research, these findings emphasise the importance of wildlife corridors between Orang NP and Kaziranga NP along the Brahmaputra River (Ojah et al. 2011, Ojah 2014). If the broader rhino landscape—the Orang NP-Laokhowa-Burhachapori WLS-Kaziranga NP complex (Kaziranga Orang Riverine Landscape), which also serves as a critical tiger corridor (Borah et al. 2010), can be preserved as a rhino metapopulation (Jhala et al. 2021), a sustainable population could be established in Laokhowa-Burhachapori WLS. Further, managing the critical grassland patches within this complex and identifying suitable habitats in this area should be prioritised. This effort could address concerns about carrying capacity (Jhala et al. 2021) and genetic erosion (Ghosh et al. 2024) by facilitating gene flow and supporting the evolutionary potential of Assam rhino populations (Ghosh et al. 2022). In addition, strong protection measures should be maintained in and around Laokhowa-Burhachapori WLS, as exploratory movements by the recolonising individuals may be vulnerable to anthropogenic threats. Protecting them should also be one of the highest immediate management priorities.

Our landscape connectivity model strongly supports the genetic connectivity patterns by identifying potential movement pathways linking Orang NP, Laokhowa-Burhachapori WLS, and Kaziranga NP. Habitat suitability in the study area was mainly influenced by NTL, NDVI, and ELE, highlighting the importance of low human disturbance, natural vegetation, and low elevation for rhino movement. The two LCPs traverse the riverine islands of the Brahmaputra, one connecting Kaziranga NP and Laokhowa-Burhachapori WLS, while the other linking the expanded areas of the 2^nd^ addition to Orang NP with the 1^st^ addition of Burhachapori WLS. The LCP between Orang NP and Laokhowa-Burhachapori WLS is shorter and has lower landscape friction, suggesting a higher potential for rhino movement from Orang NP to Laokhowa-Burhachapori WLS. While both LCPs pass through significant stretches of agricultural lands, the Orang NP-Laokhowa-Burhachapori WLS connection contains more natural vegetation, making it a more feasible movement corridor than the Kaziranga NP-Laokhowa-Burhachapori WLS link. Past land-use research also showed a ∼20% decline in vegetation cover in the Brahmaputra valley and increases in built-up and cultivated areas (∼385% and ∼8%, respectively; Debnath et al. 2022). Additionally, Areendran et al. (2020) reported that roughly 23% of the Kaziranga NP-Laokhowa-Burhachapori WLS corridor is heavily impacted by agriculture and human development.

The Brahmaputra landscape is among the last floodplain ecosystems of its kind within large river systems worldwide (Debnath et al. 2022). The expansion of protected areas connecting Kaziranga NP, Laokhowa-Burhachapori WLS, and Orang NP, along with the restoration of grassland habitats, has formed one of the largest continuous habitats, providing a vital corridor for species such as the Indian rhinoceros, Asian elephants, tigers, and wild water buffaloes. This connectivity helps sustain ecological links across the region (Borah et al. 2010; Qureshi et al. 2014; Areendran et al. 2020; Jhala et al. 2021). Jhala et al. (2021) and Ghosh et al. (2024) emphasize the conservation importance of this landscape, noting that preserving a metapopulation or natural gene flow minimizes extinction risk and maintains high genetic diversity for rhinos over time. Additionally, Laokhowa-Burhachapori WLS is designated as a priority site for rhino reintroduction under the Indian Rhino Vision (IRV) 2020, underscoring its crucial role in rhino conservation. Maintaining landscape connectivity should also be prioritized in other rhino habitats in Assam, such as between Pobitora WLS and Orang NP, which are separated by approximately 26 sq. km. of Brahmaputra floodplain, and occasional rhino movements have been recorded between these parks (Talukdar and Sarma 2007). Protecting and restoring this riverine habitat could enable natural dispersal and enhance gene flow between these two genetically distinct groups (Ghosh et al. 2024). Similarly, other critical rhino habitats, such as the forest areas between Jaldapara NP and Buxa Tiger Reserve in West Bengal, and Dudhwa NP and Katerniaghat WLS in Uttar Pradesh, also have similar potential to retain rhino metapopulations.

Our results indicate that expansion of protected areas may be insufficient unless functional connectivity is established. This allows for natural dispersal and gene flow across the landscape. The natural recolonization of Laokhowa-Burhachapori WLS demonstrates that landscape-scale conservation efforts can recover ecological processes that disappeared after local extinction. Therefore, maintaining the Kaziranga Orang Riverine Landscape as a metapopulation is essential to preserve genetic diversity, enable natural range expansion, and ensure the long-term survival of the Indian rhinoceros in Assam.

## Acknowledgements

We thank the Ministry of Environment, Forest and Climate Change (MoEF&CC), Government of India, for their invaluable support in implementing this project. Thanks are also extended to the Forest Departments of Assam for providing necessary permits and help while collecting the field samples and for providing the tissue samples. We are grateful to Shri Sonam Kumar Gupta, former range officer, Burhachapori WLS and the all the foresters of Laokhowa-Burhachapori WLS and Dr Somyita Sur and Sri Munin Gogoi from WWF India for helping with the field sampling. We thank the World Wide Fund for Nature-India (WWF-India) for their financial, field and logistic support. Our appreciation goes to Dr. Tista Ghosh for her review and suggestions while writing the manuscript. We thank Mr. Dhruv Jain, Mr. Deb Ranjan Laha and Dr. Bilal Habib, Nodal officer, NTCA Tiger Cell, Wildlife Institute of India for providing us with the boundary maps of the landscape for analyses. Finally, we thank the Director, Dean, Research Coordinator, and Nodal Officer of the Wildlife Forensics and Conservation Genetics Cell, Wildlife Institute of India for their support.

